# Methylotroph Natural Product Identification by Inverse Stable Isotopic Labeling

**DOI:** 10.1101/2021.04.26.441494

**Authors:** Dale A. Cummings, Alice I. Snelling, Aaron W. Puri

## Abstract

Natural products are an essential source of bioactive compounds. Isotopic labeling is an effective way to identify natural products that incorporate a specific precursor; however, this approach is limited by the availability of isotopically-enriched precursors. We used an inverse stable isotopic labeling approach to identify natural products by growing bacteria on a ^13^C-carbon source and then identifying ^12^C-precursor incorporation by mass spectrometry. We applied this approach to methylotrophs, ecologically important bacteria predicted to have significant yet underexplored biosynthetic potential. We demonstrate this method identifies *N*-acyl homoserine lactone quorum sensing signals produced by diverse methylotrophs grown on three different one-carbon compounds. We then apply this approach to simultaneously identify five uncharacterized signals produced by a methylotroph, and link these compounds to their synthases. We envision that this method can be used to identify other natural product classes synthesized by methylotrophs and other organisms that grow on relatively inexpensive ^13^C-carbon sources.

---

Natural products are an important source of small molecule therapeutics,^1^ agricultural compounds,^2^ and other bioactive metabolites. Precursor feeding studies are commonly used to determine the biosynthetic route to a compound of interest, and can also be used to identify previously undetected compounds produced by an organism.^3–6^ However, this approach can be limited by the synthetic or commercial availability of an isotopically-enriched precursor. Alternatively, in an inverse stable isotopic labeling approach, a fully ^13^C-labeled organism is fed a ^12^C-biosynthetic precursor to identify natural products that incorporate this precursor. This incorporation can be detected by a negative shift in the mass-to-charge ratio (*m/z*) consistent with incorporation of all or part of the ^12^C-precursor. With this approach researchers can use any precursor without having to synthesize or purchase a ^13^C-labeled version. Inverse stable isotopic labeling has been previously used to help determine the biosynthetic origins of the cofactor pyrroloquinoline quinone^7^ and an isocyanide-containing antibiotic.^8^ However, to our knowledge, this approach has not been used to systematically identify previously uncharacterized natural products.

Inverse stable isotopic labeling is particularly well-suited for bacteria that can grow on inexpensive and readily available ^13^C-carbon sources, such as methylotrophs. Methylotrophs are bacteria that grow on reduced compounds with no carbon-carbon bonds, such as methane, methanol, methylamine, and dimethyl sulfide.^9^ These organisms play important roles in biogeochemical cycling and bioremediation,^10,11^ as well as plant-microbe interactions.^12^ Genomic analysis also predicts that methylotrophs possess the biosynthetic potential to produce myriad natural products that have not yet been identified.^13^ Identifying and characterizing these molecules can help scientists understand how these important organisms interact with each other and their environment, discover new drug leads, and help synthetic biologists optimize the desirable activities of methylotrophs using exogenous small molecules.

A benefit of working with methylotrophs is that ^13^C-labeled versions of their one-carbon growth substrates are relatively inexpensive. For this reason, a methylotroph was used to create ^13^C-labeled nucleic acids from ^13^C-methanol for DNA structural studies by NMR spectroscopy.^14^ Stable isotopic labeling experiments have also been repeatedly used to identify active methylotrophs in the environment.^15,16^ We sought to take advantage of this benefit to aid in the discovery of methylotroph natural products.

As a proof of concept, we applied this method to identify quorum sensing (QS) signals produced by methylotrophs. QS is a form of chemical communication used by bacteria to regulate gene expression in a cell density-dependent manner.^17^ In one well-characterized form of QS, Gram-negative proteobacteria use *N*-acyl-homoserine lactones (acyl-HSLs) produced by LuxI-family synthases. Acyl-HSL signals vary in their acyl chain, but all possess a common homoserine lactone derived from methionine via *S*-adenosyl-*L*-methionine (Figure 1A).^18^ These signals are detected by LuxR-family receptors, which are also transcription factors that regulate gene expression upon signal binding. Methylotrophs often possess QS systems, and several strains have three or more annotated *luxI*-family synthase genes in their genome.^19,20^ The majority of these genes are predicted to encode CoA-utilizing enzymes known to produce noncanonical signals.^21^ However, relatively few methylotroph acyl-HSL signals have been characterized, and many cannot be predicted based on the amino acid sequences of their synthases.^13^

**Figure 1.**
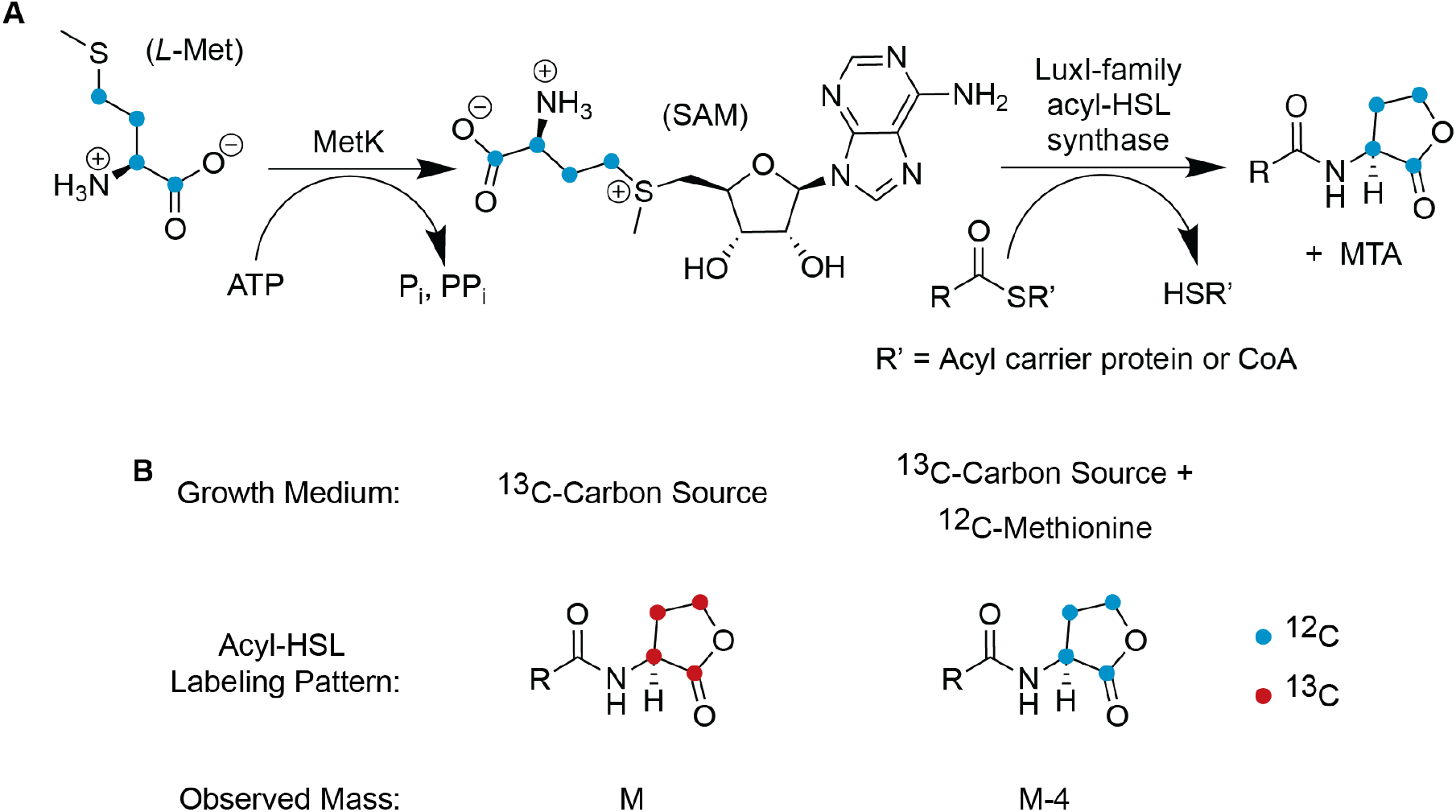
(A) Acyl-HSL biosynthesis incorporates methionine. *L*-Met: *L*-methionine. MetK: *S*-adenosyl-*L*-methionine synthase. SAM: *S*-adenosyl-*L*-methionine. MTA: 5’-methylthioadenosine. (B) The inverse stable isotope labeling approach applied to acyl-HSL signals. Note that the acyl carbonyl carbon is not denoted as ^12^C or ^13^C as it can be derived from endogenous or exogenous sources and therefore may or may not be labeled when an organism is grown on a ^13^C-carbon source.

Acyl-HSL quorum sensing signals are often detected using reporter assays, in which a strain is engineered so that the LuxR-family receptor activates the expression of a reporter gene upon signal binding.^22^ However, signal detection is dependent on the specificity of the receptor used, which can lead to false negatives. Reporter assays are often paired with thin layer chromatography to correlate detected signals with the retention factors of known standards, which can be inaccurate. Liquid chromatography coupled to tandem mass spectrometry (LC-MS/MS) can also be used to identify acyl-HSLs based on known fragmentation patterns,^23,24^ but access to these instruments may still limit routine analysis using this method, and fragmentation patterns may differ for some acyl-HSLs. For example, the aryl-HSL *p*-coumaroyl-HSL does not produce the characteristic *m/z* 102 HSL fragment.^25^

Detection of methionine incorporation into the common HSL portion of acyl-HSLs is a more generalizable method that has been successfully used to detect QS signals.^6,25 14^C-labeled methionine can be used in feeding studies to indirectly identify signals by comparing radioactivity retention via HPLC to the retention times (RTs) of known acyl-HSL standards. We sought to build off of this approach by using an inverse stable isotopic labeling method with ^12^C-methionine to identify QS signals produced by methylotrophs (Figure 1B). We show this method works with three ^13^C-labeled one-carbon sources (methane, methanol, and methylamine) in two diverse methylotrophs. We then apply the method to simultaneously identify five QS signals produced by *Methylorubrum rhodinum* DSM2163^26^ (referred to as DSM2163), including three which could not be predicted based on sequence homology to characterized QS systems. Finally, we link these signals to the synthase genes responsible for their production.

## Inverse stable isotopic labeling identifies previously characterized acyl-HSL signals

To verify the utility of the inverse stable isotopic labeling approach for identifying natural products, we began by applying it to a methylotroph that produces acyl-HSL signals that have been previously characterized. The methanol-oxidizing *Methylorubrum extorquens* PA1 produces a single LuxI-family signal synthase, MlaI, with 100% amino acid sequence identity to the characterized version found in *Methylorubrum extorquens* AM1.^19,27^ Vorholt and coworkers previously demonstrated that MlaI produces two QS signals with unsaturated acyl chains: *N*-(2-*trans*-7-*cis*-tetradecenoyl)-*L*-homoserine lactone (referred to as 2*E*,7*Z*-C_14:2_-HSL) and *N*-(7-*cis*-tetradecenoyl)-*L*-homoserine lactone (7*Z*-C_14:1_-HSL) (Figure 2).^27^

**Figure 2.**
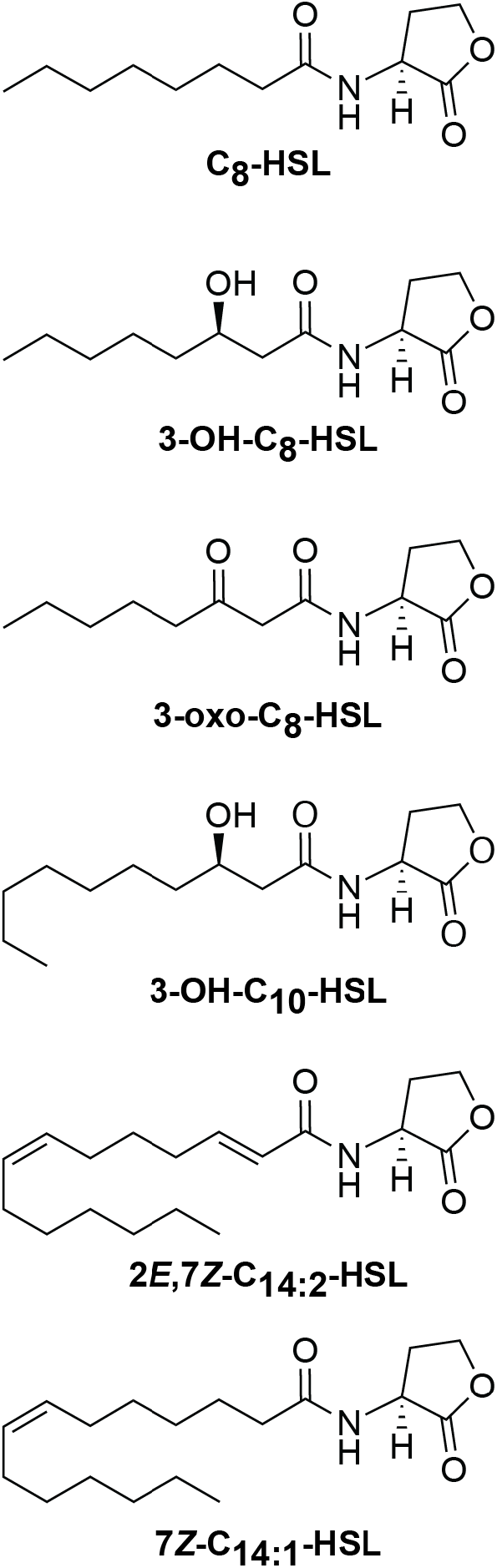
Acyl-HSLs identified in this work. Stereochemistry was not determined and is inferred based on previous studies. All biologically produced acyl-HSLs that have been characterized have homoserine lactones with an *L* stereocenter. The hydroxyls in 3-OH-C_8_-HSL and 3-OH-C_10_-HSL are shown as 3*R* because their synthases are annotated as acyl carrier protein (ACP)-linked (KEGG Orthology term K13060) and these enzymes use acyl-ACPs from fatty acid biosynthesis^30,31^ where the stereoselective β-ketoacyl acyl carrier protein reductase FabG produces 3*R*-OH fatty acyl chains.^32^

In order to streamline the inverse stable isotopic labeling procedure for future screening of diverse bacteria with potentially different growth rates, we chose to add the ^12^C-methionine precursor at the beginning of growth. We grew the *M. extorquens* PA1Δ*cel* strain CM2730 (PA1)^28^ with ^13^C-methanol and different concentrations of ^12^C-methionine. We extracted the supernatant with acidified ethyl acetate, analyzed the extract by LC-MS, and detected features using the program MZmine2.^29^ We identified two features with *m/z* values of 322 and 324, which correspond to protonated versions of 2*E*,7*Z*-C_14:2_-HSL and 7*Z*-C_14:1_-HSL, respectively, that have incorporated the four carbons of ^12^C-methionine but are otherwise fully ^13^C-labeled (Figure S1 and Table 1). When we cultured PA1 with ^12^C-methanol as a control, we identified features with the same RTs but with *m/z* values of 308 and 310, which correspond to the protonated ^12^C-versions of 2*E*,7*Z*-C_14:2_-HSL and 7*Z*-C_14:1_-HSL, respectively. Together, these results show that the inverse labeling approach can be used to correctly identify acyl-HSLs.

**Table 1.**
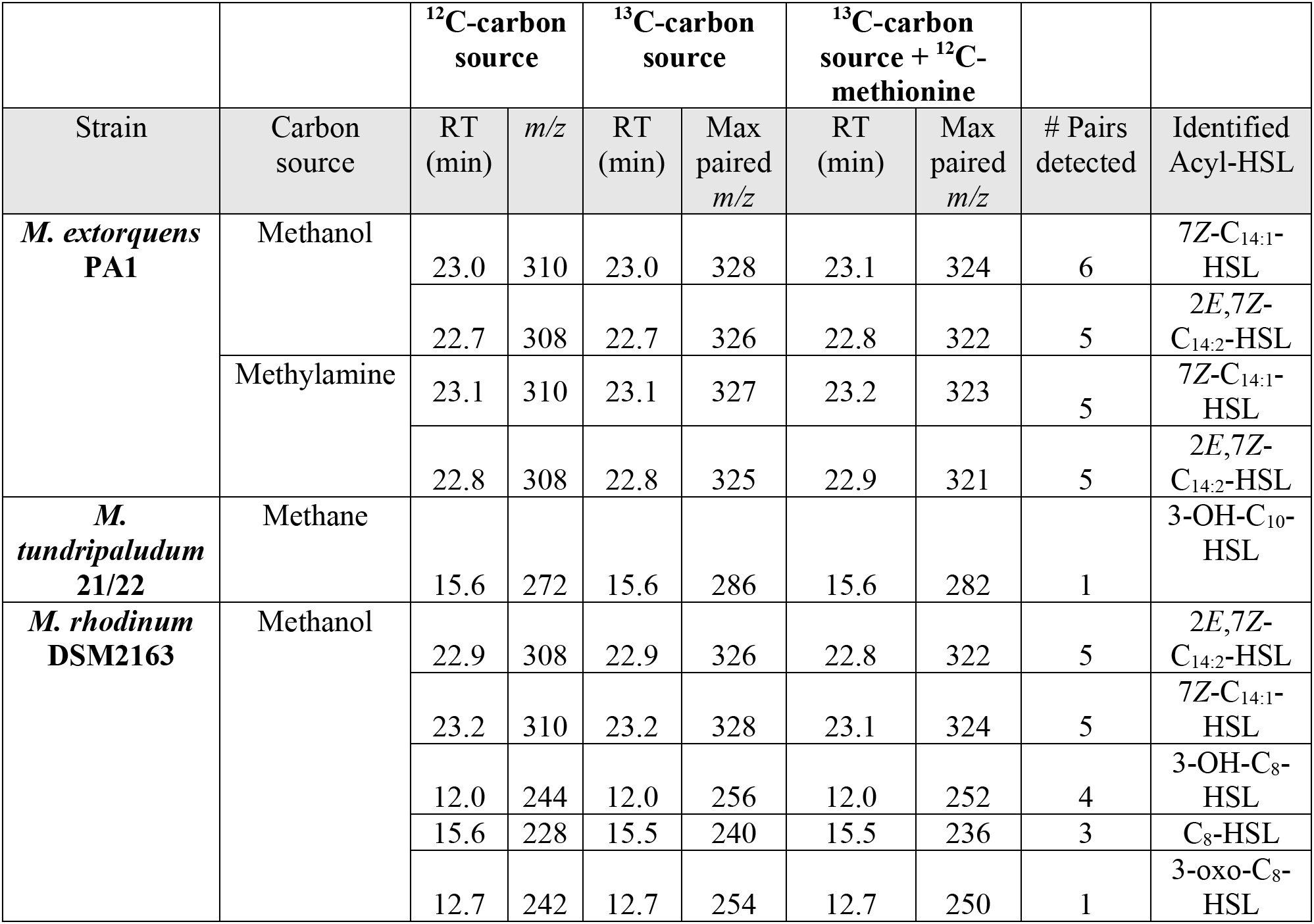
Identification of acyl-HSL signals using the inverse stable isotopic labeling approach. Max paired *m/z* refers to the maximum feature *m/z* observed in that condition where the desired *m/z* difference of four units was observed compared to the other ^13^C-condition. Note the RT differences between PA1 and DSM2163 for 2*E*,7*Z*-C_14:2_-HSL and 7*Z*-C_14:1_-HSL were resolved when smaller injection volumes were used after initial screening (see Figure S3).

The methionine titration indicated that the optimal concentration of the ^12^C-methionine precursor was 0.5 mM when added at the beginning of growth (Figure S1). Higher-than-necessary label concentrations may result in slowed growth or nonspecific incorporation of ^12^C-precursor carbons due to toxicity or methionine metabolism, respectively. We did not observe any fully ^13^C-labeled acyl-HSLs using our workflow upon addition of ^12^C-methionine at any concentration we tested. This indicates that for acyl-HSL natural products it is necessary to include a separate ^13^C-carbon source condition with no precursor as a benchmark to identify ^12^C-enriched features in the ^13^C-carbon source + ^12^C-methionine condition.

In addition to detecting fully ^13^C-labeled acyl-HSLs in the PA1 culture grown on ^13^C-methanol without addition of the ^12^C-precursor, we also detected several features with the same RT but stepwise decreases of one *m/z* unit. We also identified the same spectral pattern in the ^13^C-methanol + ^12^C-methionine condition, but decreased by four *m/z* units, corresponding to ^12^C-methionine incorporation (Figure S2). Many methylotrophs in the alphaproteobacteria class, including *M. extorquens*, assimilate reduced one-carbon compounds such as methanol using the serine cycle.^9^ In the serine cycle, half of the carbon input is derived from carbon dioxide, which we did not label in our system and could therefore result in incomplete ^13^C-labeling of metabolites of interest. The fact that we do observe fully ^13^C-labeled acyl-HSLs is likely because ^13^C-carbon dioxide is rapidly produced in the culture via central metabolism. When we grew PA1 on the carbon source ^13^C-methylamine, we did not identify complete ^13^C-labeling but this did not affect our ability to identify a decrease of four *m/z* units corresponding with ^12^C-methionine incorporation (Table 1), because the extent of acyl-HSL ^13^C-labeling was consistent between the ^13^C-methylamine and ^13^C-methylamine + ^12^C-methionine conditions.

We wrote a Python script to identify pairs of features with matching RTs and a decrease of four *m/z* units in the ^13^C-carbon source + ^12^C-methioine condition compared to the ^13^C-carbon source condition (see Methods). For the RTs corresponding to 2*E*,7*Z*-C_14:2_-HSL and 7*Z*-C_14:1_-HSL, our script identifies several pairs of features with the desired *m/z* difference due to the aforementioned incomplete labeling (Table 1). Notably, out of all the feature pairs identified in the PA1 culture extract, the two acyl-HSL signals have the most identified pairs that correspond with a single feature at the same RT in the ^12^C-carbon source control (Table S1). We can therefore use this knowledge to prioritize identified features that are more likely to be acyl-HSLs.

In order to determine if this method is more broadly applicable, we applied it to the methane-oxidizing bacterium *Methylobacter tundripaludum* 21/22 (21/22), which was previously shown to produce the QS signal *N*-(3-hydroxydecanoyl)-*L*-homoserine lactone (3-OH-C_10_-HSL) (Figure 2).^33^ When we grew 21/22 with ^13^C-methane, we identified a feature with an *m/z* value of 286, corresponding to the protonated and fully ^13^C-labeled 3-OH-C_10_-HSL. We detected a feature with the same RT and a decrease of four *m/z* units in the ^13^C-methane + ^12^C-methionine condition (Table 1), indicating ^12^C-methionine incorporation. These results show that the inverse stable isotopic labeling method can be broadly applied, as it works with the methane-oxidizing gammaproteobacterium 21/22 as well as the methanol- and methylamine-oxidizing alphaproteobacterium PA1.

## Simultaneous identification of five acyl-HSL signals produced by the methylotroph DSM2163

Next, we applied the inverse stable isotopic labeling method to identify the signals produced by the methanol-oxidizer *M. rhodinum* DSM2163 (DSM2163), which possesses three annotated LuxI-family acyl-HSL synthases. A previous study used a reporter assay to speculate which acyl-HSL signals are produced by this strain, but the exact signals were not identified.^20^ We identified several features with the signature decrease of four *m/z* units when DSM2163 was grown on ^13^C-methanol + ^12^C-methionine compared to ^13^C-methanol alone. Three of these features corresponded to the signals *N*-octanoyl-*L*-homoserine lactone (C_8_-HSL), *N*-(3-hydroxyoctanoyl)-*L*-homoserine lactone (3-OH-C_8_-HSL), and *N*-(3-oxooctanoyl)-*L*-homoserine lactone (3-oxo-C_8_-HSL) (Figure 2 and Table 1), which we confirmed by high-resolution mass spectrometry (Table S2) as well as by verifying these compounds have identical LC-MS retention times to commercial standards (Figures S3A-C).

Our method also identified features in the DSM2163 culture corresponding to the signals 2*E*,7*Z*-C_14:2_-HSL and 7*Z*-C_14:1_-HSL, and these signals had identical LC-MS retention times (Figures S3D and S3E) and high-resolution MS/MS spectra (Table S3) compared to the signals produced by PA1. This is consistent with the fact that one of the annotated DSM2163 acyl-HSL synthases shares 91% amino acid identity with MlaI from PA1. Notably, the characteristic *m/z* 102 HSL fragment was not detected in the tandem mass spectra of 2*E*,7*Z*-C_14:2_-HSL, likely because it contains an a, β unsaturated acyl chain (Table S3), and would therefore not have been identified using LC-MS/MS methods that focused on this transition. Together, these results demonstrate that the inverse labeling method can successfully identify a total of five acyl-HSL QS signals produced by the methylotroph DSM2163.

## Heterologous expression links acyl-HSL signals with their synthases

As more bacterial genomes and metagenomes are sequenced, there is an increasing demand for predicting how these organisms interact in the environment. Linking QS signal synthases with the signals they produce helps researchers make these predictions based on sequence homology. We constructed a strain for heterologous expression of *luxI*-family acyl-HSL synthase genes to link synthases with their signal products. We created an unmarked deletion strain of PA1 called AWP227 that no longer produces acyl-HSL signals by knocking out the acyl-HSL receptor gene *mlaR* and the synthase gene *mlaI*. We then used AWP227 to express each of the three DSM2136 synthase genes separately on a plasmid to determine which signals are produced by each synthase.

When we heterologously expressed the DSM2163 synthase gene Ga0373200_3300, we observed features corresponding to production of C_8_-HSL (Figure 3A) and 3-oxo-C_8_-HSL (Figure 3B) that were not present in the no-plasmid AWP227 control. Heterologous expression of Ga0373200_1920 also resulted in production of C_8_-HSL (Figure 3A), as well as 3-OH-C_8_-HSL (Figure 3C). It is therefore unclear if one or both of these synthases are responsible for the C_8_-HSL detected in the DSM2163 culture. We also confirmed that the synthase with 91% amino acid identity to MlaI (Ga0373200_956) produces the signals 2*E*,7*Z*-C_14:2_-HSL and 7*Z*-C_14:1_-HSL (Figures 3D and 3E). All gene locus tags refer to the Joint Genome Institute Integrated Microbial Genomes & Microbiomes data management system (JGI IMG/M).^34^ Vorholt and coworkers previously named a LuxI-family synthase that produces *N*-hexanoyl-*L*-homoserine lactone (C_6_-HSL) and C_8_-HSL MsaI for *<u>M</u>ethylobacterium* (now reclassified *Methylorubrum*) <u>s</u>hort-chain <u>a</u>cyl-HSLs.^27^ We therefore name Ga0373200_1920 MsaI2 and Ga0373200_3300 MsaI3 based on our results, and also refer to Ga0373200_956 as MlaI.

**Figure 3.**
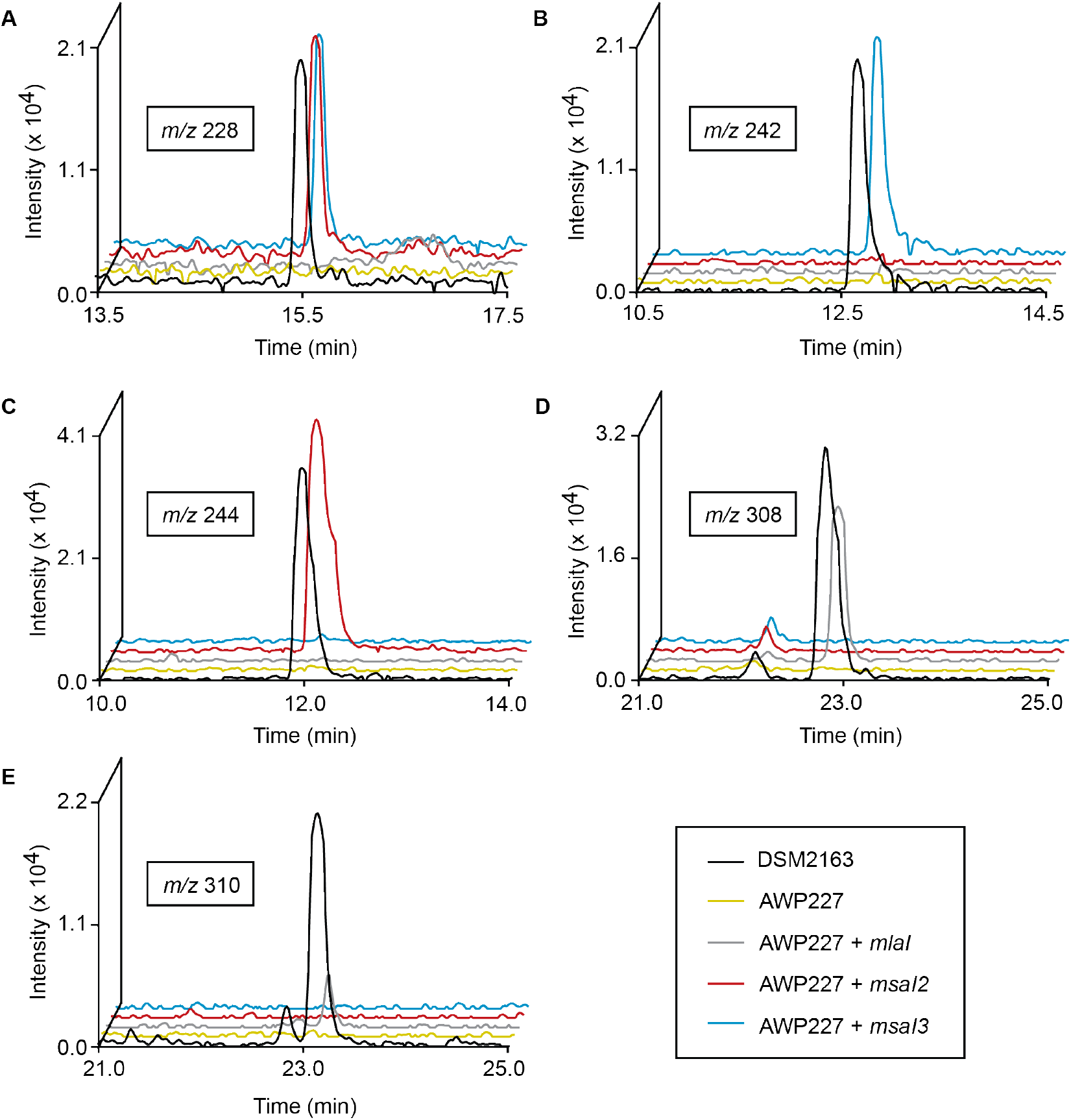
Linking *M. rhodinum* DSM2163 LuxI-family acyl-HSL synthases with their products. Extracted ion chromatograms of supernatant extracts for the listed strains for the *m/z* ranges (A) 228.0-228.5, corresponding to C_8_-HSL, (B) 242.0-242.5, corresponding to 3-oxo-C_8_-HSL, (C) 244.0-244.5, corresponding to 3-OH-C_8_-HSL, (D) 308.0-308.5, corresponding to 2*E*,7*Z*-C_14:2_-HSL, and (E) 310.0-310.5, corresponding to 7*Z*-C_14:1_-HSL.

The acyl-HSL signals produced by MsaI2 and MsaI3 could not be predicted on the basis of amino acid sequence alone. MsaI2 shares only 58% amino acid identity with RaiI from *Rhizobium etli* ISP42, which was reported to produce C_8_-HSL and 3-OH-C_8_-HSL.^35^ MsaI3 does not share >50% amino acid identity with any currently characterized LuxI-family synthases. Together, these annotations will aid in the future prediction of QS signal production from bacterial genome and metagenome sequences. Furthermore, our heterologous expression strain can be used in the future to identify QS signals produced by synthases from non-methylotrophic species using the inverse labeling approach.

Natural products are a vetted source of bioactive compounds with medicinal and agricultural value. Isotopic labeling is often used to identify and characterize natural products, however the labeled precursors required for these studies can be difficult to obtain commercially or synthetically. Here we use an inverse stable isotopic labeling approach to identify natural products via incorporation of unlabeled precursors, which significantly expands precursor availability for natural product studies. The inverse stable isotopic labeling method can be applied in the future to identify other classes of natural products based on a common precursor, including those which may not be readily available in an isotopically labeled form. Feeding studies using isotopically enriched precursors are extremely widespread, and the inverse stable isotopic labeling method is applicable in any situation where a traditional method can be used. However, this method is especially well-suited for organisms that grow on inexpensive ^13^C-carbon sources, including methanotrophs and autotrophs such as cyanobacteria, which are known to have significant biosynthetic potential.^36^

## Methods

### Key Reagents

^13^C-labeled methanol, methane, and methylamine were purchased from Cambridge Isotope Laboratories. C8-HSL was purchased from Millipore Sigma. All other acyl-HSLs were purchased from Cayman Chemical.

### Inverse labeling experiments

Exponentially growing bacterial cultures were pelleted at 16,100 rcf for one minute and resuspended in growth medium with no carbon source. Subsequently, three separate six milliliter cultures were inoculated with the resuspended strain at a starting OD of 0.02. The ^12^C-carbon source was added to one culture, the ^13^C-carbon source to the second, and the ^13^C-carbon source plus 500 nM ^12^C-methionine to the last culture. The carbon sources used were 50 mM methanol, 50 mM methylamine, or 50% (v/v) methane. Cultures were grown until reaching stationary phase (OD of approximately 0.8) and then were centrifuged at 4,800 rcf for ten minutes. The resulting supernatant was extracted twice with an equal volume of ethyl acetate containing 0.01% acetic acid, and the combined organic extract was evaporated to dryness using a nitrogen stream and stored at −20°C until analysis by LC-MS.

### LC-MS for acyl-HSL signal detection

Dried culture supernatant extracts were resuspended in 200 microliters of 1:1 water:acetonitrile, and subsequently 65 microliters were injected onto an Agilent 1260 Infinity liquid chromatography system connected to an Agilent 6120 single quadrupole mass spectrometer operating with positive polarity and a mass range of 150-1500 *m/z*. A Waters Xselect HSS T3 column (2.5 μm particle size, 2.1 mm x 50 mm) held at 30°C was used for reverse phase separation with a flow rate of 0.4 mL/min. Solvent A: Water + 0.1 % formic acid, Solvent B: Acetonitrile + 0.1% formic acid. Gradient: 0-2 min, 0% B. 2-32 min, 0-100% B. 32-35 min, 100% B. 35-36 min, 100-0% B. 36-38 min, 0% B. The limit of detection for known standards on this setup was 125 nM (Figure S4), which corresponds to a limit of approximately 5 nM in the original bacterial culture.

### LC-MS analysis

Raw data files in netCDF format were exported using Agilent OpenLab CDS (rev C.01.07). Features were detected using MZmine version 2.53^29^ using the following workflow: 1. Mass detection (centroid, noise level 1.0E3). 2. ADAP chromatogram builder^37^ (minimum group size 5 scans, group intensity threshold 1.0E3, min highest intensity 5.0E3), *m/z* tolerance 0.3). 3. Chromatogram deconvolution (local minimum search, chromatogram threshold 30%, search minimum 0.1 min, minimum relative height 10%, minimum absolute height 6.0E3, minimum ratio of peak top/edge 2, peak duration 0-2 min). 4. Adduct search (RT tolerance 0.1 min, adducts [M+Na-H] and [M+NH_3_] selected, *m/z* tolerance 0.2, max relative peak height 200%). 5. Feature list rows filter (remove identified adducts). Subsequently, isotopes were removed from ^12^C samples using the Isotopic peaks grouper (*m/z* tolerance 0.2, retention time tolerance 0.1 min, monotonic shape required, maximum charge 3, representative isotope most intense), and the three feature lists were aligned in the order ^13^C-carbon source + ^12^C-methionine, ^12^C-carbon source, ^13^C-carbon source using the Join aligner (*m/z* tolerance 0.3, weight of *m/z* 50, retention time tolerance 0.1 min, weight for retention time 50). The alignment was exported in .csv format with the row retention time as a common element and peak *m/z* as the data file element. Features containing the desired four *m/z* unit difference in the ^13^C-carbon source and ^13^C-carbon source + ^12^C-methionine samples were then detected using a custom Python script (available at https://github.com/purilab/inverse).

## Acknowledgements

This work is dedicated to Mary E. Lidstrom (University of Washington) for her 70^th^ birthday. This work was supported by National Institutes of Health grant R00 GM118762 (to AWP). We thank members of the Puri Lab for reading the manuscript and for helpful discussions. We thank Yanfen Fu (Facebook, Inc.) for useful discussions about methylotroph metabolism and isotopic labeling that led to this project. We thank N. Cecilia Martinez-Gomez (UC Berkeley) for *M. extorquens* PA1 Δ*cel* strain CM2730, and Ming Hammond (University of Utah) for generously sharing her LC-MS system and other lab equipment. We thank Daniel Petras (University of Tübingen) for sharing Python code and for helpful discussions about LC-MS analysis.

## Author Contributions

AWP and DAC designed the experiments. DAC, AIS, and AWP performed the experiments. AWP wrote the manuscript. AWP and DAC edited the manuscript. All authors read and approved of the final version of the manuscript.

