## Supplemental Information for "Methylotroph Natural Product Identification by Inverse Stable Isotopic Labeling"

**Table of Contents**


| Section | Page |
|---|---|
| Supplementary Methods | 3 |
| Supplementary Figures | 4 |
| Supplementary Tables | 8 |
| Supplementary References | 11 |

### Supplementary Methods

*Routine bacterial culturing*

Strains used in this study are listed in Table S4. *Escherichia coli* strains were grown in lysogeny broth (LB) at 37ºC. *M. extorquens* PA1 and *M. rhodinum* DSM2163 were grown at 30ºC in modified ammonium mineral salts (AMS) medium,[1] with the addition of 0.1% (m/v) yeast extract for strain DSM2163. AMS contains 0.2 g/L $MgSO_4·7H_2O$, 0.2 g/L $CaCl_2·6H_2O$, 0.5 g/L $NH_4Cl$, 30 µM $LaCl_3$, and 1X trace elements. 500X trace elements contains 1.0 g/L $Na_2$-EDTA, 2.0 g/L $FeSO_4·7H_2O$, 0.8 g/L $ZnSO_4·7H_2O$, 0.03 g/L $MnCl_2·4H_2O$, 0.03 g/L $H_3BO_3$, 0.2 g/L $CoCl_2·6H_2O$, 0.6 g/L $CuCl_2·2H_2O$, 0.02 g/L $NiCl_2·6H_2O$, and 0.05 g/L $Na_2MoO·2H_2O$. Final concentrations of 4 mM phosphate buffer pH 6.8 and 50 mM $^{12}C$- or $^{13}C$-methanol were added prior to use and cultures were shaken at 200 rpm. *M. tundripaludum* 21/22 was grown in modified nitrate mineral salts (NMS) medium,[1] which is the same as AMS with 1 g/L $KNO_3$ substituted for the $NH_4Cl$. *M. tundripaludum* 21/22 was cultured at room temperature (22-24ºC) in an atmosphere of 50% (v/v) methane in air. For routine culturing, plates were incubated in sealed jars while liquid cultures were grown in 18- by 150-mm tubes sealed with rubber stoppers and aluminum seals shaken at 200 rpm.

*Plasmid construction*

Plasmids used in this study are listed in Table S5. Primers used in this study are listed in Table S6. All plasmids were constructed using Gibson Assembly[2] and selection was performed with kanamycin (50 µg/mL) both in *E. coli* and *M. extorquens* PA1 strains.

*Genetic manipulation*

Genetic manipulation of strain PA1Δ*cel* (CM2730) and the derivative strain AWP227 was performed at 30ºC. Verified plasmids were conjugated into these strains using the *E. coli* donor S17-1.[3] 500 µL of exponentially growing cultures (OD 0.4-0.6) of the donor and recipient strains were pelleted at 16,100 rcf for one minute and resuspended in 500 µL sterile ultrapure $H_2O$. These strains were then pelleted again and the two pellets were combined in a total volume of 50 µL sterile ultrapure $H_2O$. Next, the entire mixture was spotted onto an AMS agar plate containing 50 mM methanol and 10% (v/v) nutrient broth and incubated for two days. Successful conjugants were selected on AMS plates containing kanamycin (50 µg/mL). To construct the unmarked deletion mutant AWP227, kanamycin-resistant integrants (single crossovers) were restreaked and then plated on an AMS plate containing 50 mM methanol and 1% (m/v) sucrose for counterselection. The resulting colonies were screened for double crossovers by kanamycin sensitivity and colony PCR before the final mutant was verified by Sanger sequencing.

*High-resolution mass spectrometry*

Mass spectrometry data were collected using a Waters Acquity I-class ultra-high pressure liquid chromatograph coupled to a Waters Xevo G2-S quadrupole time-of-flight mass spectrometer. An Acquity UPLC BEH C18 column (2.1 x 50 mm) was used for separation and resolving samples. Solvent A: Water + 0.1 % formic acid, Solvent B: Acetonitrile + 0.1% formic acid. The sample was eluted from the column using a ten minute linear solvent gradient: 0-0.1 min, 1% B; 0.1 - 10 min, 100% B. The solvent flow rate was 0.45 mL per minute. Mass spectra were collected in positive ion mode, with following parameters: 3 kV capillary voltage; 25 V sampling cone voltage; 150 °C source temperature; 500 °C desolvation temperature; nitrogen desolvation at 800 L/hr. The fragmentation spectra were collected using the same parameters with a 10-25 eV collision energy ramp. The lockspray solution was 200 pg/µL leucine enkephalin. The lockspray flow rate was 6 µL/min. Sodium formate was used to calibrate the mass spectrometer. The acquired mass spectra were processed using Masslynx 4.1 software.

**Supplementary Figures**

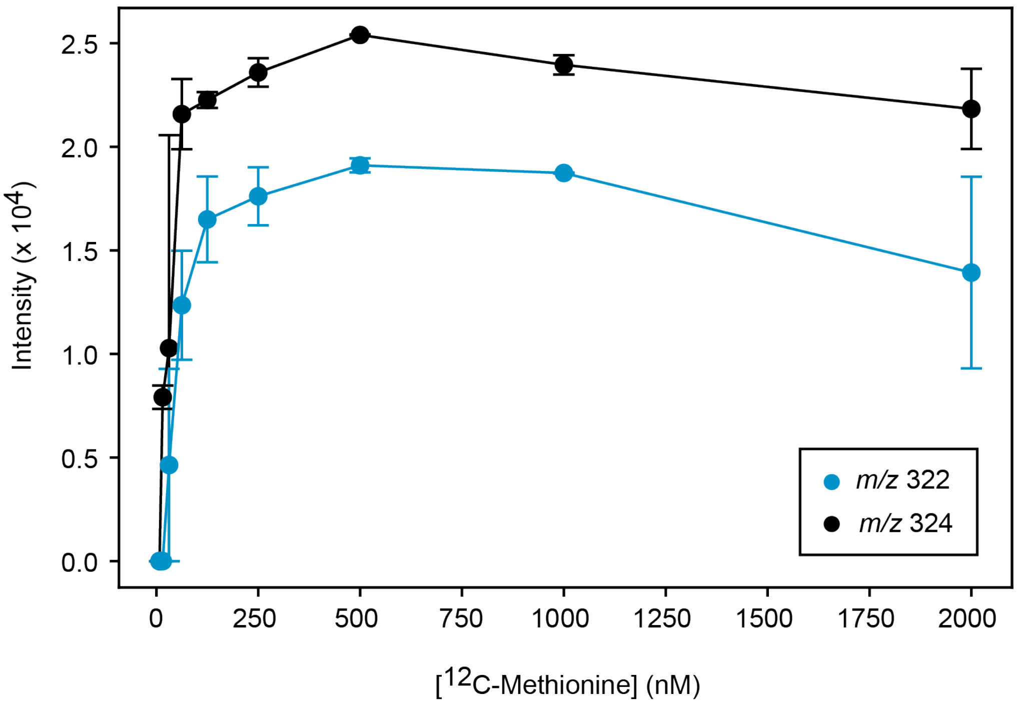


**Figure S1.** Optimization of the $^{12}$C-methionine precursor concentration. PA1 was grown in $^{13}$C-methanol with varying concentrations of $^{12}$C-methionine. At the completion of the experiment, the peak heights of features corresponding to inverse labeled 2*E*,7*Z*-$C_{14:2}$-HSL (*m/z* 322) and 7*Z*-$C_{14:1}$-HSL (*m/z* 324) were detected by LC-MS. Data represent the mean and range of two independent experiments. Peaks with intensities below 6000 were excluded during the analysis.

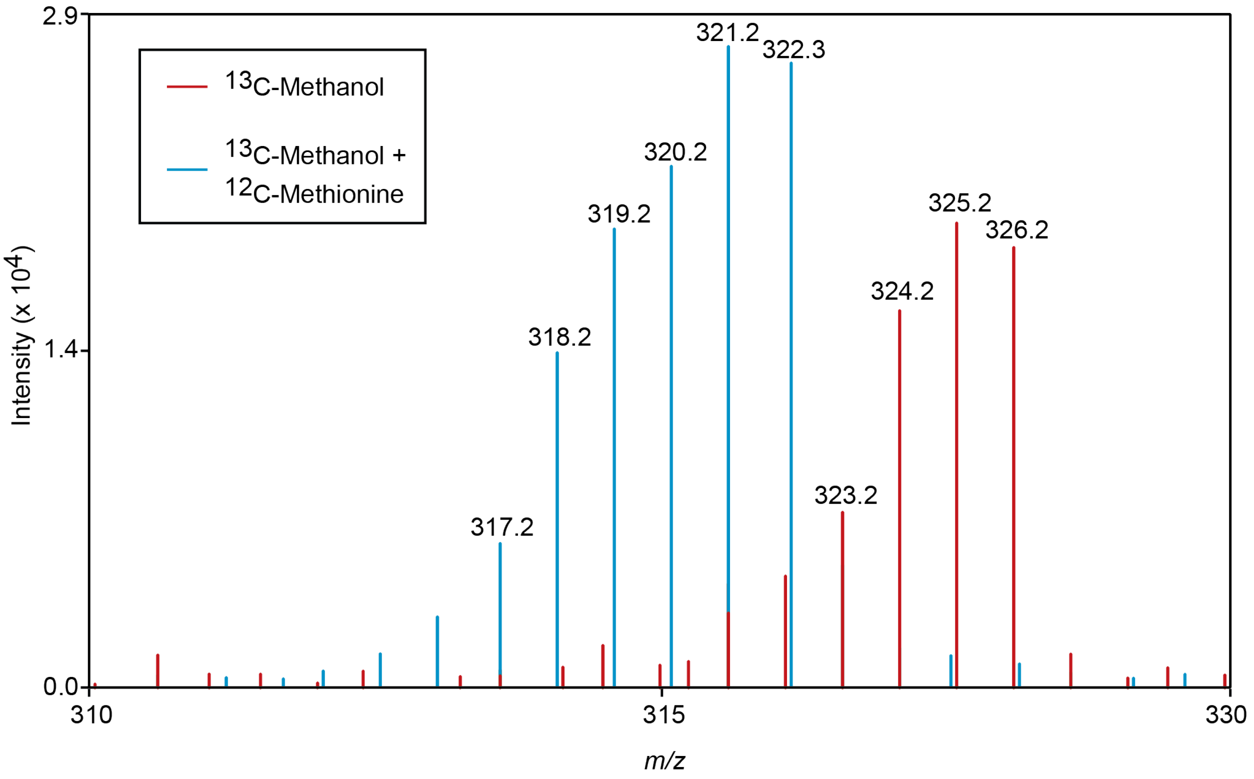


**Figure S2.** Incomplete $^{13}$C labeling of 2*E*,7*Z*-$C_{14:2}$-HSL produced by PA1 grown on $^{13}$C-methanol. Protonated and fully $^{13}$C-labeled 2*E*,7*Z*-$C_{14:2}$-HSL is *m/z* 326.

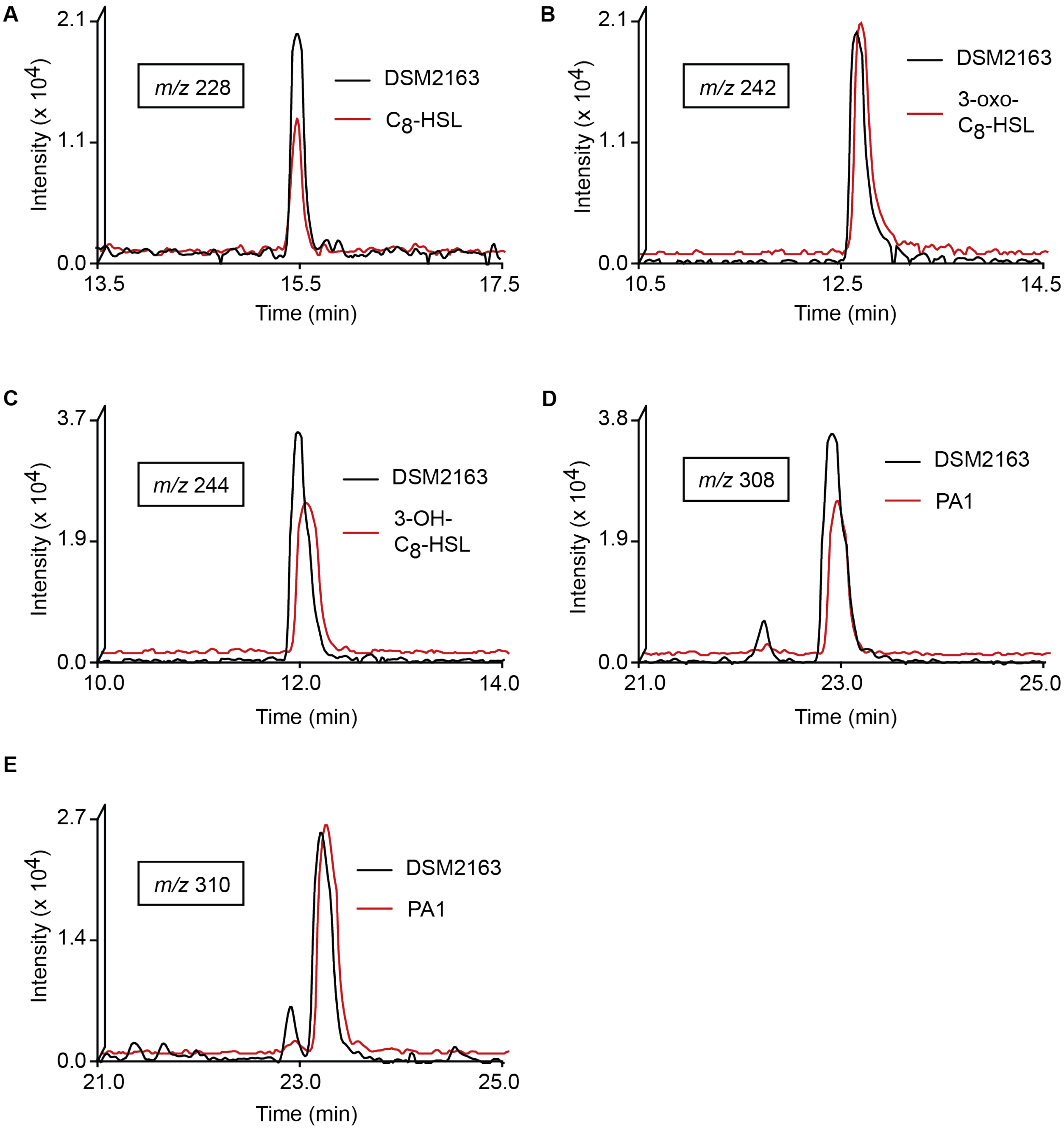


**Figure S3.** Verification of DSM2163 signal identities. Extracted ion chromatograms of commercial standards or supernatant extracts for listed strains for the *m/z* ranges (A) 228.0-228.5, corresponding to $C_8$-HSL, (B) 242.0-242.5, corresponding to 3-oxo-$C_8$-HSL, (C) 244.0-244.5, corresponding to 3-OH-$C_8$-HSL, (D) 308.0-308.5, corresponding to 2*E*,7*Z*-$C_{14:2}$-HSL, and (E) 310.0-310.5, corresponding to 7*Z*-$C_{14:1}$-HSL.

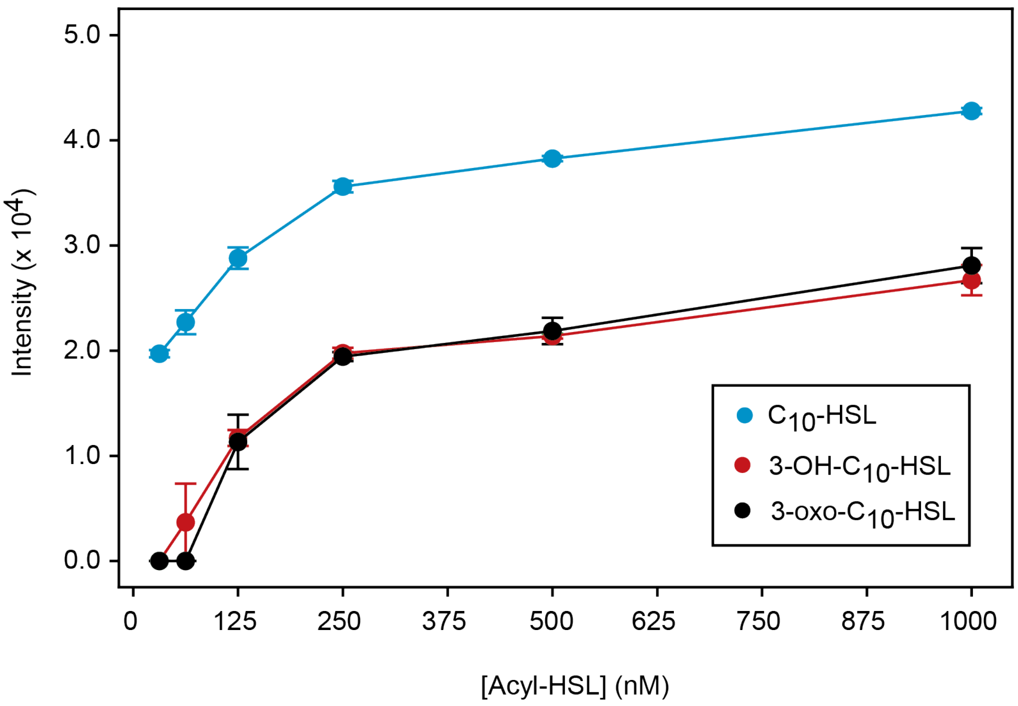


**Figure S4.** Limits of acyl-HSL detection by LC-MS. Data represent the mean and range of two independent LC-MS runs. Peaks with intensities below 6000 were excluded during the analysis.

### Supplementary Tables

**Table S1.** Output of Python script for each independent inverse stable isotopic labeling experiment. Please see attached spreadsheet.

**Table S2.** High-resolution mass spectra for identified DSM2163 signals.

| | **[M+H] Expected** | **[M+H] Observed** | **Mass error (ppm)** |
|---|---|---|---|
| $C_8$-HSL | 228.1600 | 228.1602 | 0.87657784 |
| 3-OH-$C_8$-HSL | 244.1549 | 244.1554 | 2.04788026 |
| 3-oxo-$C_8$-HSL | 242.1392 | 242.1392 | 0 |
| 7*Z*-$C_{14:1}$-HSL | 310.2382 | 310.2383 | 0.322332969 |
| 2*E*,7*Z*-$C_{14:2}$-HSL | 308.2226 | 308.2228 | 0.648881685 |

**Table S3.** LC-MS/MS spectra for unsaturated $C_{14}$-HSLs produced by PA1 and DSM2163. The top 20 most intense signals are shown for each sample.

| **2*E*,7*Z*-$C_{14:2}$-HSL** | | | | **7*Z*-$C_{14:1}$-HSL** | | | |
|---|---|---|---|---|---|---|---|
| **PA1** | | **DSM2163** | | **PA1** | | **DSM2163** | |
| ***m/z*** | **Intensity** | ***m/z*** | **Intensity** | ***m/z*** | **Intensity** | ***m/z*** | **Intensity** |
| 95.087 | 6.55E+04 | 95.0867 | 1.38E+05 | 102.0564 | 1.74E+05 | 102.0563 | 9.57E+04 |
| 207.1749 | 6.54E+04 | 207.1746 | 1.32E+05 | 310.2387 | 1.11E+05 | 310.2382 | 6.19E+04 |
| 109.1021 | 4.26E+04 | 109.102 | 8.52E+04 | 191.1801 | 5.43E+04 | 191.18 | 3.01E+04 |
| 81.072 | 3.73E+04 | 81.0716 | 7.27E+04 | 209.1908 | 5.40E+04 | 209.1902 | 2.99E+04 |
| 308.2229 | 2.95E+04 | 308.2224 | 6.42E+04 | 121.102 | 5.13E+04 | 121.102 | 2.75E+04 |
| 189.1644 | 2.24E+04 | 189.1639 | 4.68E+04 | 135.1176 | 3.89E+04 | 135.1176 | 2.02E+04 |
| 123.1177 | 2.04E+04 | 123.1172 | 4.25E+04 | 109.1023 | 3.13E+04 | 109.1023 | 1.72E+04 |
| 97.1028 | 1.81E+04 | 133.1015 | 3.66E+04 | 95.087 | 2.47E+04 | 95.0871 | 1.37E+04 |
| 125.0966 | 1.79E+04 | 97.1024 | 3.57E+04 | 311.2419 | 2.30E+04 | 311.2413 | 1.22E+04 |
| 133.1016 | 1.76E+04 | 125.0966 | 3.35E+04 | 107.0866 | 1.49E+04 | 107.0868 | 8.72E+03 |
| 107.0868 | 1.68E+04 | 107.0864 | 2.97E+04 | 93.0716 | 1.30E+04 | 93.0715 | 7.00E+03 |
| 119.0864 | 1.14E+04 | 119.086 | 2.33E+04 | 149.1329 | 1.11E+04 | 81.072 | 6.69E+03 |
| 137.0968 | 1.13E+04 | 137.0962 | 2.28E+04 | 81.0721 | 1.07E+04 | 149.133 | 6.07E+03 |
| 208.1782 | 9.78E+03 | 208.178 | 1.88E+04 | 97.1029 | 9.64E+03 | 83.0876 | 5.59E+03 |
| 83.0875 | 9.32E+03 | 93.0713 | 1.76E+04 | 83.0875 | 9.06E+03 | 97.1027 | 5.38E+03 |
| 93.0716 | 8.77E+03 | 121.1016 | 1.76E+04 | 97.0663 | 8.93E+03 | 74.0626 | 4.86E+03 |
| 121.1021 | 8.34E+03 | 179.1797 | 1.57E+04 | 123.1173 | 8.03E+03 | 97.0664 | 4.79E+03 |
| 147.1172 | 7.47E+03 | 83.0873 | 1.54E+04 | 210.1944 | 7.91E+03 | 292.2284 | 4.66E+03 |
| 179.1799 | 7.43E+03 | 147.1172 | 1.38E+04 | 292.2277 | 7.89E+03 | 103.0596 | 4.60E+03 |
| 309.2264 | 6.30E+03 | 309.2255 | 1.38E+04 | 103.0596 | 7.86E+03 | 123.1173 | 4.49E+03 |

**Table S4.** Strains used in this study.

| Strain | Puri Lab Strain Collection Number | Description | Reference |
|---|---|---|---|
| *E. coli* TOP10 | EAWP2 | F– *mcrA* Δ(*mrr-hsdRMS-mcrBC*) Φ80*lacZ*ΔM15 Δ*lacX74 recA1 araD139* Δ(*ara leu*) 7697 *galU galK rpsL* ($Str^R$) *endA1 nupG* | Invitrogen |
| *E. coli* S17-1 λpir | EAWP3 | Donor strain. $Tp^R$ $Sm^R$ *recA thi pro hsd*($r^-m^+$) RP4-2-Tc::Mu::Km Tn7 λ*pir* | 3 |
| *M. tundripaludum* 21/22 | AWP100 | Methanotroph isolated from Lake Washington (Seattle, WA, USA) sediment | 4 |
| *M. extorquens* PA1 Δ*cel* (CM2730) | AWP226 | Pink pigmented methylotroph. Δ(*celABC*-Mext_1370) | 5 |
| *M. extorquens* PA1 AWP227 | AWP227 | Pink pigmented methylotroph. Δ(*celABC*-Mext_1370) Δ*mlaRI* | This study |
| *M. rhodinum* DSM2163 | AWP225 | Pink pigmented methylotroph. | 6 |
| *M. extorquens* PA1 AWP227 + pAWP349 | AWP228 | Heterologous expression of DSM2163 *mlaI* | This study |
| *M. extorquens* PA1 AWP227 + pAWP350 | AWP229 | Heterologous expression of DSM2163 *msaI2* | This study |
| *M. extorquens* PA1 AWP227 + pAWP351 | AWP230 | Heterologous expression of DSM2163 *msaI3* | This study |

**Table S5.** Plasmids used in this study.

| Plasmid | Description | Reference |
|---|---|---|
| pAWP78 | IncP-based expression vector | 7 |
| pCM433kanT | Sucrose counterselection vector for creating clean deletion mutants | 7 |
| pAWP227 | pCM433kanT containing flanks to knock out *mlaRI* in PA1 | This study |
| pAWP349 | Expressing DSM2163 Ga0373200_956 gene under the PA1 *dnaG* (Mext_0611) promoter (400 bp upstream sequence) | This study |
| pAWP350 | Expressing DSM2163 Ga0373200_1920 gene under the PA1 *dnaG* (Mext_0611) promoter (400 bp upstream sequence) | This study |
| pAWP351 | Expressing DSM2163 Ga0373200_3300 gene under the PA1 *dnaG* (Mext_0611) promoter (400 bp upstream sequence) | This study |

**Table S6**. Primers used in this study. Homology regions used for Gibson Assembly are bolded.

| Primer Name | Sequence (5' to 3') | Description |
|---|---|---|
| oAWP186_ 433KTV1_fwd | ATGTGCAGGTTGTCGGTGTC | For amplifying the pCM433kanT backbone. oAWP186 and 160 were used to amplify one piece, and oAWP159 and 187 were used to amplify the other. |
| oAWP160_ 433KTV1_fwd | **ATAAAGGTGAATCCCATAGGGCAGGAGCTA**TAATCTCGAGTCCCGTCAAG | |
| oAWP159_433KTV2_fwd | TAGCTCCTGCCCTATGGGAT | |
| oAWP187_433KTV2_rev | TGGTAACTGTCAGACCAAGTTTACTC | |
| oAWP259_78V_fwd1 | TTGTCGGGAAGATGCGTGAT | For amplifying the pAWP78 backbone. |
| oAWP254_78V_rev1 | CAGCTCACTCAAAGGCGGTA | |
| oAWP894_227U_fwd1 | **ATATGAGTAAACTTGGTCTGACAGTTACCA**GTGATCCTGGCGACCGTCACC | For amplifying flanks to knock out PA1 *mlaRI* (Mext_4513/14) using pCM433kanT. |
| oAWP895_227U_rev1 | **TTTTGCCGGGCCATG**TTCGCCCGGCTCGCCTCCTG | |
| oAWP896_227D_fwd1 | **GGCGAGCCGGGCGAA**CATGGCCCGGCAAAATCATC | |
| oAWP897_227D_rev1 | **CGTGCATCACGACACCGACAACCTGCACAT**GCGGCATCTTGGGCATAAGAAA | |
| oAWP994_PdnaG_fwd | **ATAACCGTATTACCGCCTTTGAGTGAGCTG**ACCTCCGCGCGGCTGCCCTC | For amplifying the PA1 *dnaG* (Mext_0611) promoter (400 bp upstream sequence). |
| oAWP995_PdnaG_fwd | **AGCGCGTACTCCGTCCCCCGAACGTTGCAT**GGGGACTCTGCTGGAAGCGG | |
| oAWP1074_349I_fwd | **TCCACGCGATCCGCTTCCAGCAGAGTCCCC**ATGATCCATGTCGTGACTGC | For amplifying DSM2163 *mlaI* (Ga0373200_956). This gene was added to pAWP349 plasmid. |
| oAWP1075_349I_rev | **GAAGGATCAGATCACGCATCTTCCCGACAA**TCACGAGGCCAGCTGC | |
| oAWP1076_350I_fwd | **TCCACGCGATCCGCTTCCAGCAGAGTCCCC**ATGATCAAAATCTTCTCCGGTGCC | For amplifying DSM2163 *msaI2* (Ga0373200_1920). This gene was added to pAWP350 plasmid. |
| oAWP1077_350I_rev | **GAAGGATCAGATCACGCATCTTCCCGACAA**TCAGGCGGCCTGGCGCA | |
| oAWP1078_351I_fwd | **TCCACGCGATCCGCTTCCAGCAGAGTCCCC**ATGCAGTTGAGCACCTTGACC | For amplifying DSM2163 *msaI3* (Ga0373200_3300). This gene was added to pAWP351 plasmid. |
| oAWP1079_351I_rev | **GAAGGATCAGATCACGCATCTTCCCGACAA**TCACGCCGCCAGCTTCAACG | |
